# THE HOXD CLUSTER IS A DYNAMIC AND RESILIENT TAD BOUNDARY CONTROLLING THE SEGREGATION OF ANTAGONISTIC REGULATORY LANDSCAPES

**DOI:** 10.1101/193706

**Authors:** Eddie Rodríguez-Carballo, Lucille Lopez-Delisle, Ye Zhan, Pierre J. Fabre, Leonardo Beccari, Imane El-Idrissi, Thi Hahn Nguyen Huynh, Hakan Ozadam, Job Dekker, Denis Duboule

## Abstract

The mammalian *HoxD* cluster lies between two topologically associating domains (TADs) matching distinct, enhancer-rich regulatory landscapes. During limb development, the telomeric TAD controls the early transcription of *Hoxd* gene in forearm cells, whereas the centromeric TAD subsequently regulates more posterior *Hoxd* genes in digit cells. Therefore, the TAD boundary prevents the terminal *Hoxd13* gene to respond to forearm enhancers, thereby allowing proper limb patterning. To assess the nature and function of this CTCF-rich DNA region *in embryo,* we compared chromatin interaction profiles between proximal and distal limb bud cells isolated from mutant stocks where various parts or this boundary region were removed. The resulting progressive release in boundary effect triggered inter-TAD contacts, favored by the activity of the newly accessed enhancers. However, the boundary was highly resilient and only a 400kb large deletion including the whole gene cluster was eventually able to merge the neighboring TADs into a single structure. In this unified TAD, both proximal and distal limb enhancers nevertheless continued to work independently over a targeted transgenic reporter construct. We propose that the whole *HoxD* cluster is a dynamic TAD border and that the exact boundary position varies depending on both the transcriptional status and the developmental context.

## INTRODUCTION

In mammals, 39 *Hox* genes play critical roles in the organization and patterning of structures during development. They are found clustered at four distinct loci, *HoxA* to *HoxD* with a high level of structural organization. While all four gene clusters are activated early on during embryogenesis (Deschamps and van Nes 2005), both *HoxA* and *HoxD* clusters are subsequently re-activated during the development of the appendicular skeleton where they also participate to the building of the limbs (Dolle et al. 1989; Zakany and Duboule 2007). In the latter case, *Hoxa* and *Hoxd* genes are controlled by large regulatory landscapes flanking the gene clusters and harboring multiple enhancers (Montavon et al. 2011; Andrey et al. 2013; Berlivet et al. 2013). These regulatory landscapes were subsequently found to coincide with topologically associating domains (TADs) (Dixon et al. 2012; Nora et al. 2012), which are defined as genome regions in which chromatin interactions occur more frequently. Such domains tend to be constitutive (Dixon et al. 2012) and hence they are mostly conserved between tissues and amongst various vertebrate species (e.g. (Woltering et al. 2014). In addition, TADs correlate with lamina associated domains (LADs) and DNA replication domains and may thus be considered as units of chromosome organization (see (Gonzalez-Sandoval and Gasser 2016)).

The *HoxD* gene cluster lies at the border between two such chromatin domains and various subsets of *Hoxd* genes respond to either limb regulatory landscape. Initially, the telomeric TAD (T-DOM), located in 3’ of the gene cluster, is active and controls the transcription of *Hoxd3* to *Hoxd11* into the most proximal part of the future limb, the arm and the forearm. Subsequently, in distal limb bud cells, T-DOM is switched off while the opposite 5’-located TAD (C-DOM) becomes active to control the expression of *Hoxd13* to *Hoxd9* into presumptive digit cells (Andrey et al. 2013; Beccari et al. 2016). Therefore, to successive waves of transcription occur, triggered by distinct enhancer landscapes and in phase with the building of the two main pieces of the future limbs.

The existence of both this switch in regulations and a strong boundary effect introduces a discontinuity in the transcription of these genes, which allows the formation of a zone of low *Hoxd* expression thus giving rise to the wrist or the ankle (Villavicencio-Lorini et al. 2010; Woltering and Duboule 2010). To produce these critical articulations, it is thus essential that enhancers located in either TADs do not regulate all *Hoxd* genes at once, which would lead to uninterrupted expression domains. Also, it was proposed that both the *Hoxd12* and *Hoxd13* products exerted a dominant negative effect over other HOX proteins (van der Hoeven et al. 1996; Zakany et al. 2004), referred to as ‘posterior prevalence’ (see references in (Duboule and Morata 1994)(Yekta et al. 2008). This strong inter-TAD border may thus exist in response to the need for *Hoxd13* and *Hoxd12* not to respond to more ‘proximal’ enhancers, since such an ectopic expression would lead to deleterious morphological effects (e.g. (Herault et al. 1997)) similar to other instances where TAD boundaries were reported to prevent ectopic interactions potentially causing diseases (Lupianez et al. 2015; Fabre et al. 2017).

The exact nature of TAD borders as well as their causality is often difficult to establish. These DNA regions are enriched in bound CTCF and cohesin subunits suggesting architectural constraints such as helping either to trigger or to prevent interactions between promoters and enhancers (Kagey et al. 2010; Sofueva et al. 2013; Zuin et al. 2014). They were shown to function in the constitutive organization of TADs (Dixon et al. 2012; Phillips-Cremins et al. 2013; Rao et al. 2014), since removal of either CTCF or of the cohesin complex affects TAD stability (Nora et al. 2017; Rao et al. 2017; Haarhuis et al. 2017; Schwarzer et al. 2017). In the case of *HoxD*, the TAD border can be mapped in the ‘posterior’ part of the cluster, between *Hoxd11* and *Hoxd12*, i.e. in a genomic region showing one of the highest GC content genome-wide and displaying nine bound CTCF sites within a 40kb large region (Soshnikova et al. 2010), as well as close to ten active promoters. In this particular genomic context, a functional dissection of this TAD border would require multiple and separate genetic interventions in-*cis* to disconnect promoter sequences from those involved in constitutive contacts and thus reveal whether enhancer-promoter contacts either impose a TAD structure, or instead are constrained by such a chromatin domain, which would form independently from any transcriptional activity.

Here we address this conundrum by analyzing *in embryo* the structural and functional effects of a series of nested deletions involving either part of the boundary region or larger pieces of the *HoxD* locus including it. We used both proximal and distal micro-dissected limb bud cells, i.e. two highly related cell types but where only one or the other of the two TADs is transcriptionally active. While small deletions elicited minor and mostly local effects, larger deletions triggered the re-arrangement of interactions leading to major chromatin reorganization. Altogether, the boundary activity for long-range contacts was surprisingly resilient and only the absence of a 400 kb large DNA region including the *HoxD* cluster itself generated a single large TAD, made out of the fusion between both T-DOM and C-DOM. We conclude that several elements in the *HoxD* locus cooperate to impose the requested segregation between the two opposite regulatory influences. The exact positioning of this boundary within the gene cluster, as well as its strength in preventing ectopic interactions may have been a powerful evolutionary cursor in the shaping of various tetrapod limb morphologies.

## RESULTS

### A TAD border within the *HoxD* cluster

In order to get insights into TAD organization around the *HoxD* locus during limb bud development, we performed Hi-C on micro-dissected distal and proximal limb bud cells isolated from E12.5 embryos. At this stage, T-DOM enhancers regulate *Hoxd* gene expression in proximal cells and are silent in distal cells, whereas C-DOM enhancers control *Hoxd* gene targets in future digit cells while they are silent in proximal cells. Therefore, the two TADs are either transcriptionally active or inactive in an exclusive manner, in the two tissue samples (Fig. 1A and B, top schemes). In both cases, the Hi-C profiles positioned the *HoxD* cluster right in-between the TADs, similar to what was initially reported either in ES cells (Dixon et al. 2012) or in CH12 lymphoblastic cells (data extracted from (Rao et al. 2014) (Supplemental Fig. S1A-D). Although the distribution of contacts was quite similar in the two cell populations, the internal organization of interactions within the TADs displayed few distinctive features at 40kb resolution. Our analysis of the CH12 lymphoblasts ENCODE datasets did not reveal any long-distance contact between the *HoxD* cluster and potential enhancer regions as both gene deserts appeared globally devoid of H3K27ac marks (Supplemental Fig S1C, bottom panel).

**Figure 1.**
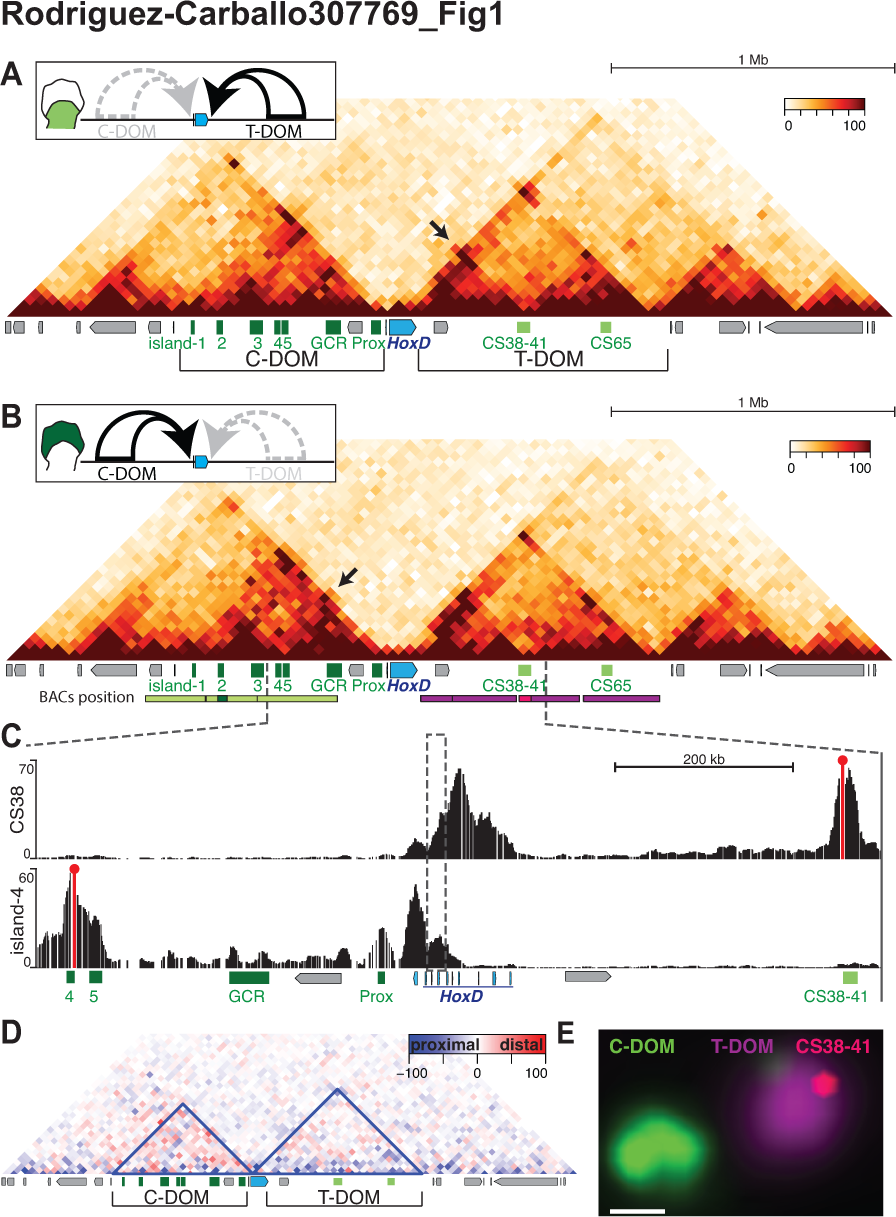
Three-dimensional organization of the *HoxD* locus in limb buds. **(A).** Hi-C heatmap using proximal E12.5 murine limb bud cells. 3Mb of chromosome 2 (mm10, 73,320,000-76,480,000) are covered. The scheme on top indicates that T-DOM is active and C-DOM inactive. The positions of the *HoxD* cluster (blue) and surrounding enhancers (green) are shown below with CS38-41 and CS65 within T-DOM, whereas Prox, GCR and islands −1 to −5 are located within C-DOM. Other surrounding genes are depicted as grey boxes. The arrow indicates contacts established between central *Hoxd* genes and CS38-41. **(B)**. Hi-C heatmap similar to **(A)** but using distal cells where C-DOM is active (top). The arrow indicates contacts between 5’-located and central *Hoxd* genes and island-3. The red and green bars below illustrate the BAC clones used in DNA-FISH experiments. **(C)**. 4C-seq tracks showing contacts established either by CS38 (top, red line), or by island-4 (bottom, red line) by using distal limb bud cells. The dashed vertical rectangle marks the boundary region. **(D)**. Subtraction of Hi-C matrices shown in **(A)** and **(B)** with distal cells in red and proximal cells in blue. The blue line demarcates the extension of the identified TADs in distal cells. **(E)**. DNA-FISH using distal limb bud cells and the series of BAC clones shown in **(B)** (scale: 500nm). The position of CS38-41 inside T-DOM is shown in red by using a fosmid clone.

In distal cells, specific contacts were established between posterior *Hoxd* genes (*Hoxd13* to *Hoxd10*) and previously defined regulatory sequences within C-DOM (islands −1 to −5, GCR and Prox (Gonzalez et al. 2007)(Montavon et al. 2011) (Fig. 1B). In proximal cells however, some of these contacts were not detected as strongly (Fig. 1A) and C-DOM showed lower contact intensities than in distal cells (p-value = 0.018), as revealed by performing a subtraction of both Hi-C datasets (Fig. 1D and Supplemental Fig. S1I). Altogether however, the two interaction maps were quite similar to one another. Likewise, T-DOM displayed only few changes in interactions when distal and proximal cells were compared (p-value = 0.87) (Fig. 1D and Supplemental Fig. S1I).

Differences between distal and proximal limb cells were nevertheless observed around the so-called CS38-41 region, which also displayed bound CTCF molecules (see below) and appeared as being itself both a boundary between the two sub-TADs found within T-DOM (Andrey et al. 2013) and a strong region of interaction with the *HoxD* cluster in the two cell populations. This region contains enhancers for limbs, caecum and mammary glands as well as a bidirectional transcription start site for the *hotdog* and *twin of hotdog* lncRNAs (Delpretti et al. 2013; Schep et al. 2016). CS38-41 was specifically contacted by the central part of the *HoxD* cluster in proximal cells only (Fig. 1A, black arrow), since in distal cells, the same region of *HoxD* interacted with the opposite C-DOM (Fig. 1B, black arrow). The Hi-C data also revealed a highly interacting region extending from the gene cluster up to CS38-41 in distal cells where it was covered by H3K27me3 marks (Andrey et al. 2013). Altogether however, no obvious interactions were detected between the two opposite regulatory landscapes. DNA-FISH analysis using independent BACs labelling either C-DOM, T-DOM or region CS38-41 (Fig. 1E, green, purple and pink, respectively), confirmed the isolated spatial conformation of both TADs and their status as independent regulatory units (see (Fabre et al. 2015)).

While these Hi-C analyses illustrated the strict partitioning between the two TADs, their resolution (40kb) made it difficult to precisely define the position of the TAD border within the *HoxD* cluster. We applied to our embryonic limb datasets various algorithms based on isolation potential (Crane et al. 2015; Shin et al. 2016) to identify these limits. This approach revealed a boundary with a dynamic position within a ca. 50 kb large DNA interval, with a more centromeric position in proximal cells and a more telomeric position in distal cells (Supplemental Fig. S1E-H, red lines). When the TopDom algorithm was applied to either murine ES cells (Dixon et al. 2012) or CH12 cells (Rao et al. 2014) datasets, a shift in the TAD border along the *HoxD* cluster was also scored. In ES cells, a micro-domain was detected involving most of the gene cluster (Supplemental Fig. S1B) (Noordermeer et al. 2014; Kundu et al. 2017) as likely associated with the presence of H3K27me3 modifications found throughout *Hoxd* genes in these cells (Bernstein et al. 2005). In CH12 however, the algorithm placed the boundary at the position of *Hoxd9* engulfing the highly active *Hoxd4* gene into T-DOM (Supplemental Fig. S1D). Therefore, the domain boundary was found located at different positions in the *Hox* cluster depending on the cell population and its set of transcribed *Hoxd* genes.

To more precisely define this TAD boundary in our experimental contexts, we used 4C-seq, an approach with a resolution below 5kb. For example, when the C-DOM island-4 was used as bait in distal limb cells, the strongest interactions were scored with the *Hoxd13* to *Evx2* region, with substantial contacts also observed over *Hoxd11* up to *Hoxd10* (Fig. 1C and Supplemental Fig. S1). Likewise, when the T-DOM located bait CS38 was used in the same cells, strong interactions were scored over *Hoxd8* and *Hoxd9* with a striking decrease in contacts over the *Hoxd10* to *Hoxd11* region (Fig. 1C and Supplemental Fig. S1G) thus positioning a border at around *Hoxd10*, whereas this border was positioned over *Hoxd11* to *Hoxd12* when the CS38 bait was used in proximal cells (Supplemental Fig. S1H and (Andrey et al. 2013). The use of these two opposite baits showed that the precise location of the boundary changed in relation with the on-off transcriptional activity of the TADs.

### Different sub-groups of transcribed *Hoxd* genes are bordered by bound CTCF and Cohesin

TAD borders are often enriched both in CpG islands and in sites bound by architectural proteins, which may be instrumental in either their formation or their maintenance (Guelen et al. 2008; Dixon et al. 2012). For instance, CTCF and the cohesin complex can form loops between distant regions and hence favor the segregation of chromatin interaction patterns (see (Phillips-Cremins et al. 2013; Hnisz et al. 2016)). The *HoxD* cluster displays a dense distribution of at least 21 identified CpG islands and contains more than ten different promoters, including coding and non-coding genes (Fig. 2A).

**Figure 2.**
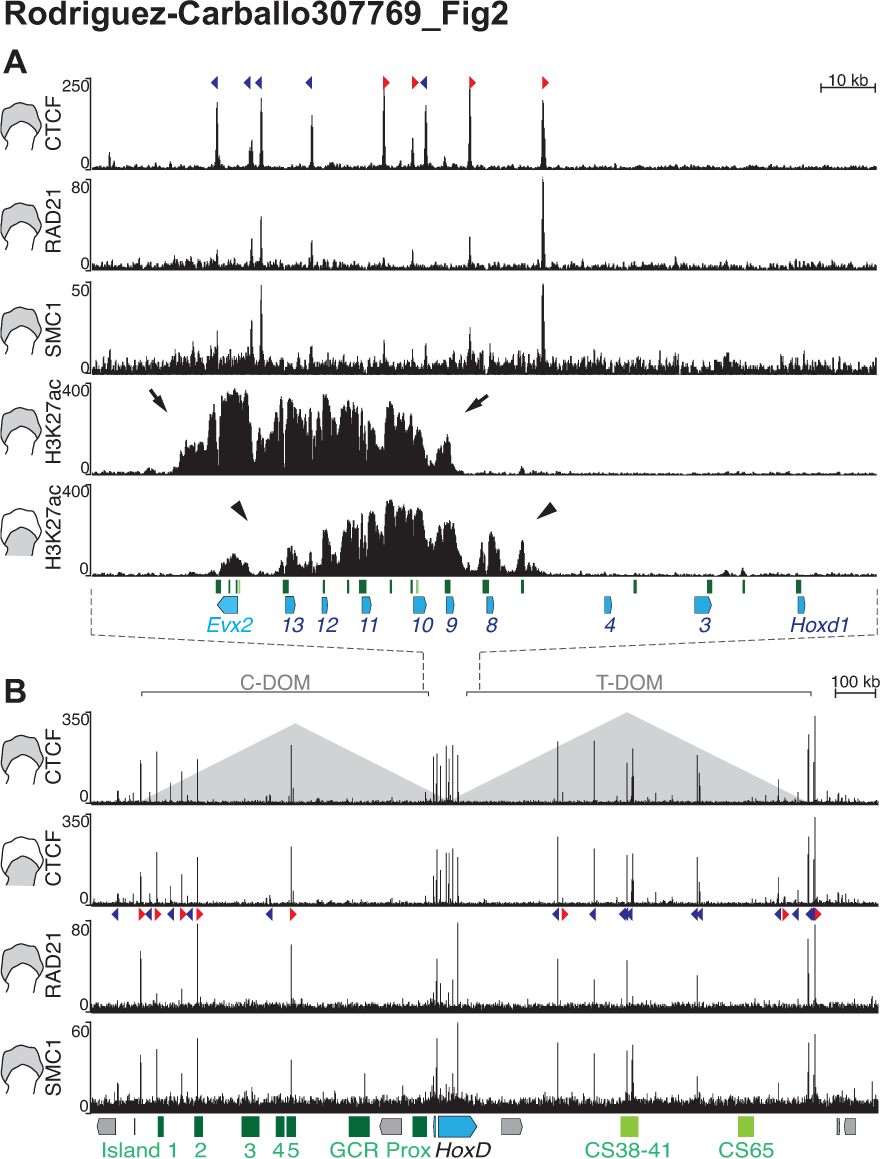
Subsets of *Hoxd* genes responding to C-DOM or T-DOM enhancers coincide with bound CTCF and cohesin complex. **(A)**. CTCF, RAD21 and SMC1 ChIP-seq profiles at and around the *HoxD* boundary region. The CTCF profile in distal cells (top track) is identical to that in proximal cells with peaks spanning the centromeric half of the gene cluster. CTCF motif orientation is shown with arrowheads (red for telomere-oriented CTCF and blue for motifs oriented towards the centromere). The profiles of RAD21 and SMC1 tend to label the extremities of the H3K27ac domains (bottom). These active domains are restricted within a large DNA interval where bound CTCF molecules are observed (arrows for distal limb and arrowheads for proximal limb). The green boxes below represent CpG islands. Diagrams on the left show which of distal or proximal cells were used. **(B)**. CTCF and SMC1 profiles along both C-DOM and T-DOM TADs (schematized as pyramids). CTCF peaks are conserved in proximal and distal cells. CTCF motif orientation is as in A. Below is the *HoxD* cluster (blue) and various regulatory elements.

In order to study the binding profile of architectural proteins over the *HoxD* locus and associated TADs, we performed ChIP experiments to identify sites bound either by CTCF in distal and proximal limb cells, or by the cohesin RAD21 and SMC1 subunits in distal limb bud cells. Noteworthy, the bound CTCF sites were mostly distributed within the centromeric half of the cluster, precisely where different blocks of genes were active in both limb cell populations, thus matching the genomic window where the boundary had been mapped (Fig. 2A). We first used MACS2 peak calling followed by consensus motif identification to classify the bound CTCF sites according to their orientations (http://insulatordb.uthsc.edu/), given that CTCF sites with divergent orientations are present at many TAD borders (Rao et al. 2014; de Wit et al. 2015; Guo et al. 2015; Tang et al. 2015; Vietri Rudan et al. 2015). Within *HoxD*, all four CTCF sites located at the centromeric side were oriented towards C-DOM whereas all those but one located at more telomeric positions faced T-DOM (Fig. 2A; colored arrowheads), suggesting an inversion in orientations between *Hoxd12* and *Hoxd11*, i.e. on either side of the TAD border observed in proximal cells.

While the sites of bound cohesin subunits mostly coincided with sequences also bound by CTCF, these subunits were enriched on both sides of the series of bound CTCF, i.e. either between *Hoxd4* and *Hoxd8* or in the *Hoxd13* to *Evx2* intergenic region. Of note, the extension of H3K27ac domains, a histone modification associated with active gene transcription, identified the distinct sub-groups of *Hoxd* genes actively transcribed either in proximal or in distal limb cells. In both cases, CTCF and cohesin were bound at‐ or in the vicinity of- both extremities of these domains (Fig. 2A), as if these proteins were used to somehow label those large target DNA regions successively accessible first by T-DOM and then by C-DOM enhancers. Bound CTCF and cohesin subunits were also scored within C-DOM and T-DOM, in particular at important regulatory sequences such as the CS38-41 region as well as at islands-1, −2 and −5, which were enriched for both CTCF and RAD21 (Fig. 2B). While most of these CTCF sites were orientated towards the *HoxD* cluster, their occupancy remained globally unchanged in the different limb cell populations (Fig. 2B), similar to the situation within the gene cluster, suggesting that CTCF alone may not bring any tissue-specificity to these regulations (Fig. 2A, B).

### Serial deletions of the TAD boundary or parts thereof

Our Hi-C and 4C-seq datasets thus located the TAD border region somewhere between *Hoxd8* and *Hoxd13*, with some variation depending on the cell type considered. To try to assess the various components of this boundary, we used a set of deletion alleles where distinct portions of this DNA interval had been removed (see (Tschopp and Duboule 2014) (supplemental Fig. S2). We used 4C-seq to document the interaction profiles generated by two opposite viewpoints located at each side of the TAD border (Fig. 3, orange bars). The *Evx2* bait lies immediately near *Hoxd13* on the centromeric side of the boundary, whereas *Hoxd4* is the first gene located clearly outside of this boundary interval on the telomeric side. Consequently, under wildtype conditions, *Hoxd4* is only expressed in proximal limb cells under the control of T-DOM, while C-DOM enhancers control *Evx2* transcripts in distal cells exclusively.

**Figure 3.**
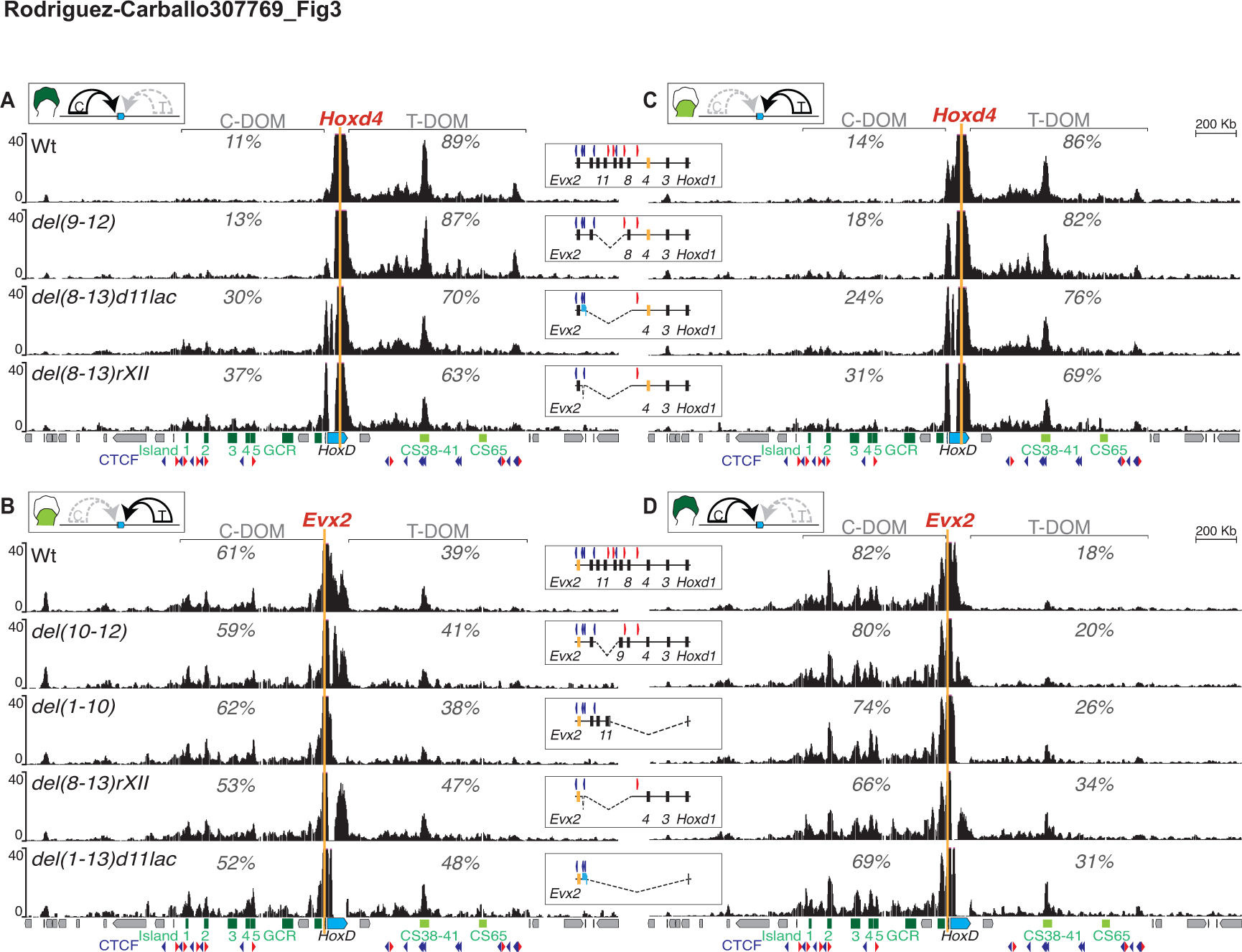
Partial deletions of the inter-TAD border. Interactions established by *Hoxd4* and *Evx2* in a set of deletion alleles including part of the boundary region. **(A)**. 4C-seq profiles by using *Hoxd4* as a viewpoint (orange line) in distal limb cells where C-DOM is active (top scheme). The control (wt), *HoxD^del(9-12)^*, *HoxD^del(8-13)d11lac^* and *HoxD^del(8-13)rXII^* are shown from top to bottom, with a schematic on the right indicating the deletion and the viewpoint (orange rectangle). The percentages reflect the ratios of contacts scored in either TADs, after excluding reads mapping to the cluster itself. Below are the gene cluster (blue) and regulatory sequences (green). **(B)**. 4C-seq profiles by using *Evx2* as a viewpoint (orange line) in proximal limb cells where T-DOM is active (top scheme). The control (wt), *HoxD^del(10-12)^*, *HoxD^del(1-10)^*, *HoxD^del(8-13)rXII^* and *HoxD^del(1-13)d11lac^* are shown from top to bottom, with a schematic on the left indicating the deletion and the viewpoint (red rectangle). **(C).** 4C-seq as in **(A)** using *Hoxd4* as a viewpoint in proximal cells where T-DOM is active. **(D).** 4C-seq profiles as in **(B)** using *Evx2* as a viewpoint in distal cells where C-DOM is active. For all panels, percentages are as for **(A)**.

We scored the interactions of these two baits in both deletion and control alleles. For each bait, we used cells where the operating TAD was on the other side of the border. In this way, we looked for ectopic gains of contacts crossing the boundary region towards a TAD containing enhancers functionally at work. We first analyzed the interactions of *Hoxd4* in distal cells, i.e. when C-DOM is fully active and T-DOM is switched off. In this situation, the control allele revealed only 11% of contacts between *Hoxd4* and C-DOM, while most of the contacts remained within T-DOM, illustrating the robustness of the boundary. A fair part of the border interval was removed in the *HoxD^del(9-12)^* allele, where the DNA region from *Hoxd9* to *Hoxd12* had been deleted. Nevertheless, very little effect if any was scored and *Hoxd4* did not appear to have increased interactions with the active C-DOM (Fig. 3A). When the larger *HoxD^del(8-13)d11lacZ^* deletion was used, where almost the full boundary region is removed and replaced by a *Hoxd11lacZ* transgene, ectopic interactions between *Hoxd4* and C-DOM started to significantly increase, from 11% to 30% of the contacts (Fig. 3A). Interactions with C-DOM increased to almost 40% when both the *Hoxd11lacZ* transgene and a small region containing a CTCF site between *Hoxd13* and *Evx2* (Supplemental Fig. S2) were further removed from this deletion (*HoxD^del(8-13)rXII^*). Even in this case, however, contacts established by the *Hoxd4* bait were still biased towards T-DOM (Fig. 3A), indicating that some boundary activity was left, perhaps associated with the few CTCF and cohesin binding sites still present on either sides of the latter deletion breakpoints (Supplemental Fig. S3A, C).

The situation was comparable, yet slightly different, when *Evx2* was used as bait. In wild type proximal limb cells where T-DOM was active and C-DOM inactive, *Evx2* already established substantial interactions with sequences located in the opposite T-DOM (Fig. 3B, 39%). Small deletions like *HoxD^del(10-12)^* or larger deletions affecting mostly genes on the telomeric side of the cluster (for example *HoxD^del(1-10)^*) did not induce any significant increase of interactions with T-DOM (Fig. 3B). *Evx2* did nevertheless increase its interactions with T-DOM whenever the more centromeric *Hoxd* genes were removed, for instance in the *HoxD^del(8-13)rXII^* allele or when the entire gene cluster was substituted by a *LacZ* reporter transgene, either in *HoxD^del(1-13)d11lac^* or in *HoxD^del(1-13)d9lac^* (Fig. 3B). In these mutant alleles, the boundary appeared slightly more efficient in blocking telomeric enhancers to access C-DOM promoters, than centromeric enhancers to leak over T-DOM genes, an observation perhaps related to the orientation of the remaining CTCF sites (see discussion). Altogether, these results suggested that the boundary was a multipartite structure, resilient to the deletion of its parts.

Ectopic interactions established by *Hoxd4* when parts of the boundary region were deleted were only marginally affected by the activity of the TADs. In proximal cells indeed, where T-DOM was active, the increased interactions of *Hoxd4* towards C-DOM in the various deletions were globally comparable to the situation in distal cells, when T-DOM was inactive (Fig. 3C). Likewise, when the same mutant alleles were compared such as *HoxD^del(8-13)rXII^*, the ectopic interactions established by *Evx2* in proximal cells were not drastically different from those scored in distal cells where C-DOM was active (Fig. 3B, D), in particular considering that the contacts between *Evx2* and T-DOM were already higher in wild type proximal cells than those between *Hoxd4* and C-DOM in distal cells (Fig 3A, B). This illustrated again that a C-DOM-located promoter was more easily attracted by the opposite T-DOM than was a T-DOM-located gene by the activity of C-DOM. This feature was also apparent when using *Hoxd13* as bait either in the wild type chromosome or on a set of deletions. In these various cases, ectopic interactions towards T-DOM were generally higher than with *Hoxd4* in the opposite situation. In addition, these interactions were increased whenever the T-DOM was transcriptionally active rather than inactive (Supplemental Fig. S4A, B).

### Ectopic inter-TADs contacts are specific and productive

We next asked whether the re-allocation of interactions observed when using some of these deletion alleles were merely structural or, alternatively, if they could elicit a transcriptional outcome. We monitored the expression of both *Hoxd4* and *Evx2* in these various alleles and observed ectopic transcriptional activation concurrent with new interactions. For example, in the *HoxD^del(8-13)rXII^* deletion, *Hoxd4* was strongly expressed in distal cells and *Evx2* in proximal cells, a situation never observed in control animals (Fig. 4A and Supplemental Fig. S5; arrowheads). Expectedly, *Evx2* transcripts were also gained in proximal cells after the deletion of the entire *HoxD* cluster (Supplemental Fig. S5B, E). Ectopic transcription precisely correlated with the re-allocation of interactions with enhancers. The quantifications of these interactions on specific regions known to be required for transcription of *Hoxd* genes in distal cells (e.g. island-2) showed that the increases in contacts were significant only in those alleles where ectopic expression was scored (Fig. 4B and Supplemental Fig. S5F).

**Figure 4.**
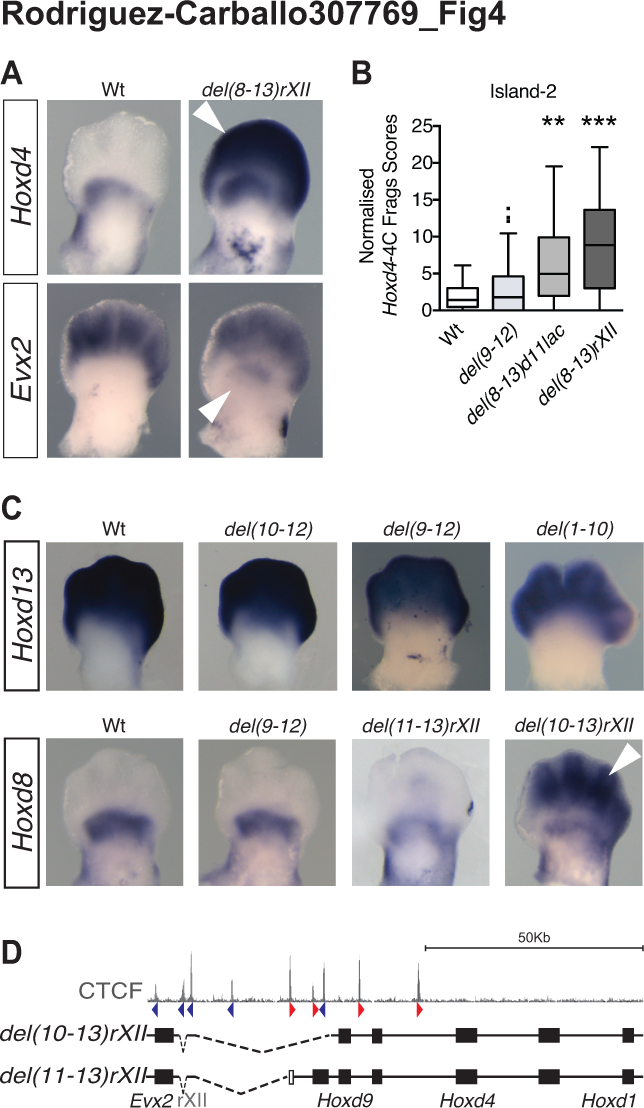
Ectopic contacts are transcriptionally productive. **(A)**. Whole mount *in-situ* hybridization (WISH) using *Hoxd4* or *Evx2* probes in E12.5 wt and *HoxD^del(8-13)rXII^* mutant forelimb buds. In this deletion allele, *Hoxd4* is massively gained in distal cells (top, arrowhead) and *Evx2* significantly gained in proximal cells (bottom, arrowhead). **(B).** Quantification of 4C-seq contacts mapping to the digit-specific island-2 enhancer in the *HoxD^del(9-12)^*, *HoxD^del(8-13)d11lac^ HoxD^del(8-13)rXII^* mutant alleles. Kruskal-Wallis test **p<0.01, ***p<0.001. **(C).** WISH using probes for *Hoxd13* and *Hoxd8* in limb buds of control and several deletion alleles, as indicated on the top, illustrating the resilience of the inter-TADs insulation effect and the gain of *Hoxd8* expression in the *HoxD^del(10-13)rXII^* (arrowhead), much weaker in the shorter *HoxD^del(11-13)rXII^*. **(D).** Schematic showing the two latter deletion alleles along with the profile of bound CTCF on top.

We confirmed these observations by analyzing the steady-state levels of *Hoxd8* mRNAs in various deletion alleles. As for *Hoxd4*, *Hoxd8* transcription remained unchanged in the *HoxD^del(9-12)^* mutant limb buds, while a weak but significant ectopic expression was scored in distal cells of E12.5 embryos carrying the *HoxD^del(11-13rXII)^* allele. Of note, *Hoxd8* expression was strongly gained in distal limbs of *HoxD^del(10-13)rXII^* mutant embryos (Fig. 4C, D; arrowhead), suggesting that the sequential removal of gene promoters and/or CTCF binding sites progressively weakened the TAD boundary (Narendra et al. 2015). However, we did not observe any ectopic expression of *Hoxd13* in proximal cells, even when a large portion of the boundary region had been removed. It is possible that the deletion was not sufficient to induce the ectopic activation of *Hoxd13,* even when interactions were gained along T-DOM such as in the *HoxD^del(1-10)^* allele (Fig. 4C and Supplemental Figs. S4C and S5C and (Zakany et al. 2004)). Altogether, neither ectopic interactions, nor the gains in transcription observed in our series of deletions could be explained by the mere change in relative position of a given *Hoxd* target gene with respect to the appropriate enhancer sequences. Because of this lack of a simple correlation, we conclude that some specific regions inside this large boundary interval are stronger than others in exerting their isolation potential.

### Deletions of the TAD boundary

In these 4C-seq experiments, both the *Hoxd4* and *Evx2* baits are located close to the deletion breakpoints and may thus be influenced by proximity effects. Consequently, while they illustrate the accessibility of target promoters to remote enhancers localized in the opposite TAD, they are not appropriate to assess the potential of the *HoxD* cluster to block inter-TADs contacts. In the latter case, ectopic interactions between enhancers located in one TAD and sequences located within the other would represent a major reorganization in local chromatin architecture. We thus performed 4C-seq using as viewpoints two regions with enhancer properties, which also seem to act as major interaction points between the *HoxD* cluster and each flanking TAD. Island-4 belongs to C-DOM and is an enhancer region strongly contacted by *Hoxd* genes transcribed in distal cells. It is not contacted in brain cells where *Hoxd* genes are inactive (Montavon et al. 2011). In contrast, the CS38 bait belongs to the CS38-41 region of T-DOM, a conserved region with multiple enhancer activities in the intestinal caecum, limbs and mammary buds (Delpretti et al. 2013; Schep et al. 2016; Beccari et al. 2016). Of note, this region contains three occupied CTCF sites, all oriented towards the cluster and is also enriched in cohesin (Fig. 2B).

These remote viewpoints confirmed that the smallest deletions containing parts of the *HoxD* TAD border did not detectably affect its insulation potential. In the *HoxD^del(9-12)^* allele for example, neither CS38 nor island-4 gained any substantial contact with the opposite TAD in either distal (Figs. 5A, B) or proximal limb bud cells (Figs. 5C, D). Moderate gains of inter-TAD interactions were nevertheless observed when larger deletions were used such as the *HoxD^del(8-13)rXII^, HoxD^del(1-10)^* or *HoxD^del(1-13)d9lac^* alleles. With island-4, relative increases of up to 10 % of interactions were scored on the opposite TAD when using small deletions, with only a weak effect associated with the on-off transcriptional status of the TAD. The gain in interactions detected between CS38 and C-DOM sequences was more significant in distal cells where C-DOM was active, than in proximal cells (Fig. 5A, C). To more precisely evaluate these effects, we generated *in silico* genomes corresponding to every deletion allele (Supplemental Fig. S2). In this way, we could analyze the cumulative signals along 3Mb around the viewpoints (Fig. 5E, F) and cluster the results according to the Euclidean distance between the curves. We noticed a clear effect related to the size of the deletions, with small deletions clustering with the control allele whereas larger deletions clustered together (Fig. 5G, H).

**Figure 5.**
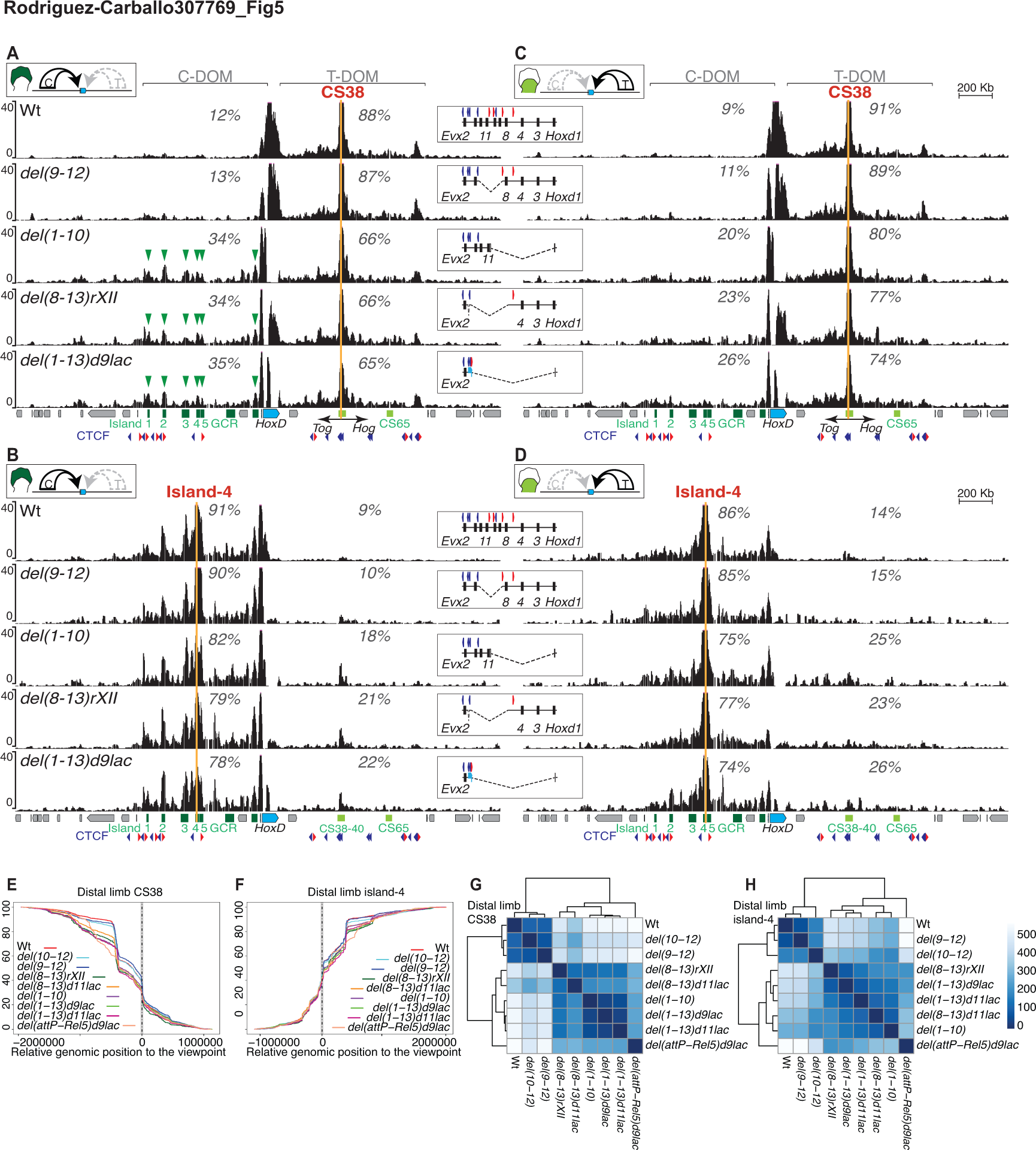
Inter-TADs contacts following partial boundary deletions. **(A).** 4C-seq interaction profiles using region CS38 as a viewpoint (orange line) in distal limb bud cells (scheme on top) of wt (n=5), *HoxD^del(9-12)^*, *HoxD^del(1-10)^*, *HoxD^del(8-13)rXII^* and *HoxD^del(1-13)d9lac^* deleted alleles (top down). Quantification of contacts in T-DOM and C-DOM expressed as percentages are as for figure 3. Schematics of the deleted region alleles are shown on the right. The *HoxD* cluster (blue) and regulatory regions (green) are depicted below, as well as the bidirectional transcription start site of the lncRNAs *Hog* and *Tog* (arrows), close to CS38. **(B).** 4C-seq interaction profiles using island-4 as a viewpoint (orange line) in distal limb bud cells (scheme on top) using the same deleted alleles as for panel **(A)**. **(C, D)**. 4C-seq profiles of CS38 and island-4 viewpoints as in **(A, B)** but in proximal cells. **(E, F)**. Cumulative sums of 4C-seq reads relative to the distance to either CS38 **(E)** or island-4 **(F)** used as viewpoints, in E12.5 distal limb cells. Colours represent different mutant alleles and the positions of viewpoints are shown by vertical dashed lines. **(G, H)** Heatmap of euclidean distances between each pair of curves obtained from panels **(E)** and **(F)**. A Ward clustering was performed on the resulting matrix.

Noteworthy, all these moderate but significant gains in interactions observed with the larger deletion alleles involved contacts with active enhancer sequences. In the *HoxD^del(8-13)rXII^, HoxD^del(1-10)^* or *HoxD^del(1-13)d9lac^* alleles for instance, CS38 contacts were gained with the islands −1 to −5, as well as with the Prox sequence in distal cells (Fig. 5A, arrowheads), whereas the contacts were not as specific in proximal cells where these enhancers are inactive (Fig. 5B). As for the *Hoxd4* bait (see Figs. 3 and 4), we asked whether such ectopic interactions could be productive and trigger transcription of T-DOM sequences into distal limb cells, an expression specificity normally excluded from this TAD (Beccari et al. 2016). We used as a readout the two lncRNAs *Hotdog (Hog)* and *twin of Hotdog (Tog)*, which are transcribed in opposite directions starting from the CS38 region (Delpretti et al. 2013). As expected from their genomic localization within T-DOM, both *Hog* and *Tog* were transcribed in control proximal limb bud cells (Supplemental Fig. S6A). In addition, both WISH and qPCR revealed a gain of *Hog* and *Tog* transcripts in distal cells dissected from all mutant embryos carrying a deleted allele where ectopic contacts with the C-DOM digit islands were scored (Supplemental Fig. S6A, B). These gains in *Hog* and *Tog* transcripts in distal cells correlated with the quantification of CS38 interactions with known distal enhancers (Supplemental Fig. S6C). However, the newly established contacts between CS38 and the C-DOM island-2 were not reflected by any substantial change in the spatial distance between these regions, as shown by DNA-FISH using the *HoxD^del(8-13)rXII^* allele (Supplemental Fig. S6D, E).

The ectopic interactions observed between the C-DOM sequence island-4 and T-DOM in the larger deletions were also slightly different depending on the activity status of each TAD. When T-DOM was inactive, in distal cells, most of the ectopic contacts involved the CS38-41 region (Fig. 5B). When T-DOM was active, in proximal cells, ectopic interactions between island-4 and T-DOM were more widespread, involving CS38-41 but also other surrounding sequences (Fig. 5D). The functional outcome, if any, of these ectopic contacts between island-4 and T-DOM sequences could nevertheless not be assessed due to the absence of any known transcription unit mapping to the C-DOM regulatory islands, which could have been used as a readout similar to *Hog* and *Tog* for T-DOM. Altogether, despite some substantial ectopic interactions observed with baits CS38 and island-4, a strong insulation between the two TADs was still observed even when the largest deletions were considered, emphasizing again the robustness of this border and its resistance to perturbations.

### Reorganization of TADs

To better document the resilience of this TAD border after large deletions, we performed Hi-C with cells where the entire gene cluster was deleted and replaced by a *Hoxd9/lacZ* reporter transgene. In our 4C analysis, this *HoxD^del(1-13)d9lac^* allele still displayed a boundary effect, even though significant inter-TADs contacts were detected (Fig. 5 and Supplemental Fig. S6). The Hi-C data highlighted this increase of inter-TAD interactions, in particular between the T-DOM sub-TAD flanking the deletion breakpoint and the most centromeric region of C-DOM (islands-1, −2, close to the *Atf2* gene), when compared to control cells (Fig. 6A-D). Despite these *de novo* interactions, a *HoxD* TAD border was called, even though it appeared less strong than in control animals as judged by using the TopDom algorithm (Shin et al. 2016) (Fig. 6A-D, dashed lines and profiles on the top). Interestingly, the *Hoxd9/lacZ* transgene contained two occupied CTCF sites with opposite orientations. While one of these sites was equally occupied at the wild type *Hoxd9* locus, the second one was only very weakly bound in the wild type condition but strongly re-enforced in the transgene present in this allele (Supplemental Figs. S2 and S3B).

**Figure 6.**
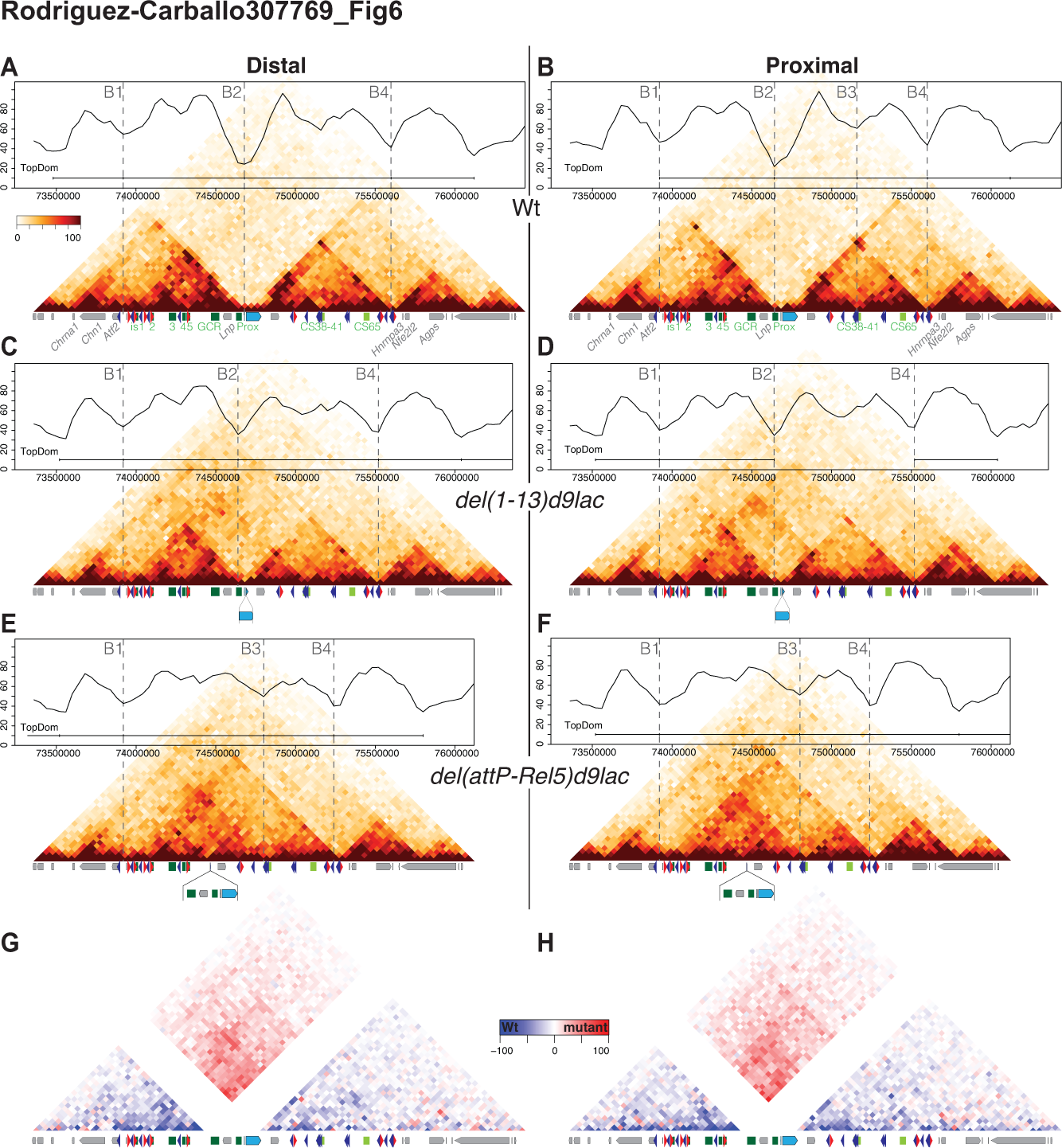
Re-organization of TADs after deletion of the *HoxD* cluster. **(A-F).** Hi-C profiles covering 3Mb of mouse chromosome 2 and centred at the *HoxD* locus (blue rectangle) in limb bud cells. The control allele (wt) is on top (**A-B**), followed by the *HoxD^del(1-13)d9lac^* (**C-D**) and *HoxD^del(attP-Rel5)d9lac^* (**E-F**) deletion alleles. For each allele, distal cells are on the left and proximal on the right. On top of the Hi-C profiles are graphs showing isolation potential based on the TopDom algorithm. A horizontal bar defines the ‘consensus’ TADs and vertical dashed lines label boundaries called by the algorithm. These boundaries are referred to as B1 to B4 for better comparison between the various alleles. B2 is the TAD boundary at the *HoxD* locus (**A-B**), which weakens in the *HoxD^del(1-13)d9lac^* allele (**C-D**) to disappear in the *HoxD^del(attP-Rel5)d9lac^* allele (**E-F**). In the latter allele, a new B3 boundary appears in distal cells (**E-F**). (**G-H**). Subtraction of Hi-C values between the *HoxD^del(attP-Rel5)d9lac^* and wild type alleles using the distal and proximal limb datasets. The mutant datasets were mapped on the wild type genome prior to subtraction.

Besides this weakened boundary, some re-organizations in intra-TAD contacts were also detected. In distal cells, C-DOM showed less heterogeneity in interactions in the mutant allele, likely due to a drastic reduction either in the number of target promoters or in CTCF sites (Fig. 6C, D and Supplemental Fig. S3B). The same was true for T-DOM, whose overall interaction density was also reduced in the mutant chromosome. In contrast, TADs located outside C-DOM and T-DOM remained unchanged (Fig. 6A-D). In proximal cells, these changes were even more pronounced. For instance, the algorithm did not detect the boundary between the two sub-TADs of T-DOM, at the position of CS38-41 (Fig. 6B, dashed line), which was routinely scored in the control allele, likely due to the reduced interactions between CS38-41 and the target promoters leading to a lower discrimination between these two sub-TADs (Fig. 6D).

In this *HoxD^del(1-13)d9lac^* allele, the *Evx2* and *Lunapark* (*Lnp*) promoters were retained, as well as the *Hoxd9* promoter present on the *Hoxd9/LacZ* transgene containing two bound CTCF sites and two to three additional CTCF sites located over *Evx2*. The persistence of these CTCF sites may account for the weak but clear boundary effect remaining between the two TADs. To clarify this issue, we used a larger deletion removing both *Evx2* and *Lnp*, in addition to the *HoxD* cluster. In this *HoxD^del(attP-Rel5)d9lac^* allele, where a ca. 400 kb large DNA segment is lacking, only the *Hoxd9/lacZ* reporter gene is left with its two bound CTCF sites, with opposite orientations. In this case, despite the presence of the two divergent CTCF sites (Supplemental Fig. S3B), the boundary disappeared and a new merged TAD formed (Fig. 6E, F).

However, the TAD formed *de novo* did not result from the fusion between the remains of C-DOM and T-DOM (from B1 to B4). Instead, it comprised the remains of C-DOM (from B1), including islands-1 to −5 and the centromeric sub-TAD (to B3) of T-DOM with a much weaker contribution of the telomeric sub-TAD of T-DOM. This was materialized by a boundary call between the newly formed TAD and the telomeric sub-TAD (from B3 to B4) in both tissues, using the same algorithm and threshold as before (Shin et al. 2016) (Fig. 6E, F). In this case, the contacts established between region 38-41 in former T-DOM and islands-1 and 2 in former C-DOM to build the new TAD coincided with the presence of clusters of bound CTCF sites in convergent orientations (Fig. 2B), which normally interact with the series of bound CTCF and cohesin found around the target *Hoxd* genes on either sides of the native *HoxD* boundary.

In both distal and proximal cells, the density of interactions within this newly formed TAD (from B1 to B3) was nevertheless below that observed in control C-DOM (B1-B2) and T-DOM (B2-B4) (Fig. 6, compare E, F with A, B), indicating that the global solidity of TAD architecture was dependent upon the presence of strong contacts points at either sides of the border. Presumably, this loss of strength in intrinsic interactions also translated into the establishment of contacts with the next telomeric boundary region, leading to the inclusion of this new TAD into a larger yet weaker structure delimited by the two original borders (B1 and B4). This marked the centromeric and telomeric extremities of the two TADs containing all remote enhancers operating at the *HoxD* locus (Fig. 6E, F). These changes were clearly detected when a subtraction was performed between the mutant and the control datasets (Fig. 6G, H).

We used DNA-FISH to see whether such a fusion between the two TADs was accompanied by a reduction in the distance between two BACs covering T-DOM and C-DOM (Fabre et al. 2015). In distal limb cells, the *HoxD^del(attP-Rel5)d9lac^* allele indeed showed a significant reduction in inter-TAD distance, when compared to control limb cells. However, this reduction was not scored when mutant proximal cells were used, further indicating that the transcriptional status of a given TAD may impact upon some of its general properties (Supplemental Fig. S7A). This difference in inter-TAD distance between mutant distal and proximal cells was not anticipated from the Hi-C dataset. This tendency was nevertheless supported by an extensive 4C analysis of this large deletion allele. For instance, when the CS38 sequence (in T-DOM) was used as bait, cross-contacts in particular with islands-1 and −2 were more noticeable in mutant distal cells than in proximal cells (Supplemental Fig. S7B, C) in agreement with the higher frequency of ‘short distances’ observed in distal mutant cells in the DNA-FISH experiment.

In this large deletion allele, the global re-organization of TAD architecture at the *HoxD* locus did not severely impact upon the neighboring TADs. On the telomeric side, the small domain including the *Hnrnpa3*, *Nfe2l2* and *Agps* genes was not affected at all (Fig. 6A-F). On the centromeric side, some contacts scored in control limbs between either *Hoxd13* or islands-1 and −2 and a sub-TAD containing the *Chn1* and *Chrna1* loci were no longer observed in the *HoxD^del(attP-Rel5)d9lac^* allele. However, here again, the interaction profiles around these transcription units were not dramatically perturbed by the important modifications occurring in the neighboring C-DOM (Fig. 6E, F).

### A recomposed enhancers landscape

In the *HoxD^del(attP-Rel5)d9lac^* allele, both T-DOM and C-DOM specific enhancers are now located within the same TAD. This is in marked contrast with the normal situation where a strict partitioning was observed between the C-DOM and T-DOM regulatory landscapes. The grouping of forearm enhancers in one TAD and of digit enhancers in the other TAD was considered as the basis of the collinear transcriptional mechanism driving *Hoxd* genes during limb development (Andrey et al. 2013). Therefore, we evaluated the impact of the fusion between TADs and the resulting promiscuity of both types of enhancers in the *HoxD^del(attP-Rel5)d9lac^* allele by using the *Hoxd9/lacZ* transgene as a readout. In this configuration, a single *Hox* promoter-*lacZ* gene is left in the center of the newly produced TAD along with proximal enhancers located in 3’ and digits regulatory islands 1 to 5 located in 5’ (Fig. 7).

**Figure 7.**
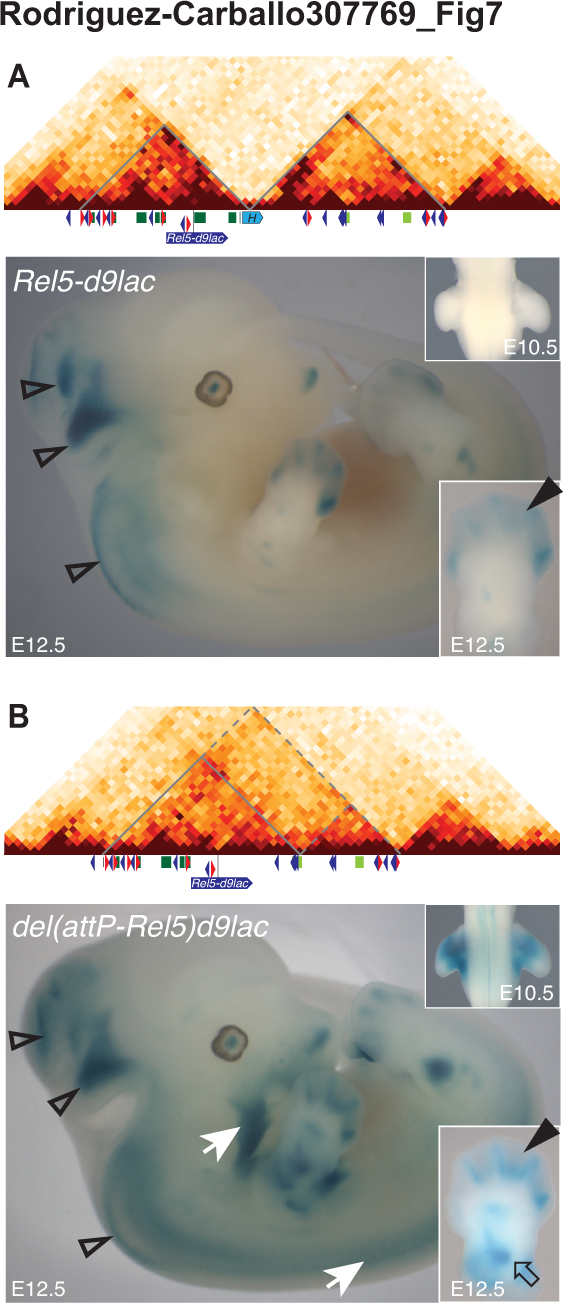
Impact of TAD merging upon enhancers specificities. **(A).** Beta-gal staining of both *HoxD^Rel5-d9lac^* (top) and (**B**) *HoxD^del(attP-Rel5)d9lac^* (bottom) mutant E12.5 embryos. The former allele is used as a non-deleted control. In addition to the staining observed in the central nervous system (open arrowheads) and in distal limbs in both cases (bold arrowheads in bottom right boxes), the large deletion shows staining emanating from T-DOM based enhancers, as exemplified in proximal limbs (open arrow in bottom right box) or in crest cells and paraxial mesoderm (white arrows). The abnormal promiscuity between enhancers does not severely impair their modes of operation. (**B**). In the deleted allele, the transgene is expressed in limb buds as early as E10.5, whereas it is not detected in the *HoxD^Rel5-d9lac^* allele (top right boxes), i.e. before deletion.

As control, we used the exact same *Hoxd9LacZ* transgene simply inserted at the *rel5* position (Spitz et al. 2003), without any deleted DNA (Fig. 7A). Because the *rel5* site is located within C-DOM, *lacZ* staining was expectedly detected in distal limb buds as well as in a column of interneurons and some part of the developing brain specific for *Evx2* regulation (Kmita 2002) (Fig. 7A, B). In the *HoxD^del(attP-Rel5)d9lac^* allele, these expression specificities were all maintained. In addition, *LacZ* expression was scored in proximal limbs, in the whisker pads as well as in a population of crest cells migrating towards the future mandibles and the axial mesoderm (Fig. 7B, white arrows), which are all expression specificities controlled by enhancers located in T-DOM (Spitz et al. 2001). Therefore, the physical separation of enhancers into two distinct TADs may not be a prerequisite for various C-DOM and T-DOM enhancers to be properly operational in space and time.

## DISCUSSION

### Alternating long-range regulations

During limb bud development, T-DOM initially drives the early phase of *Hoxd* gene activation, whereas C-DOM subsequently regulates the second wave of transcription. The boundary between these two TADs is dynamic and more or less well defined. In ES cells, in the absence of transcription, the entire *HoxD* gene cluster forms a dense domain, which is positioned at the border between the two TADs (Noordermeer et al. 2011; Dixon et al. 2012; Williamson et al. 2014; Kundu et al. 2017) (Fabre et al. 2015). In the different developing tissues analyzed thus far, however, the position of the TAD border matches the transcriptionally active *versus* inactive transition in the gene cluster, reflecting the preferential interaction of transcribed genes with the active TAD (e.g.(Andrey et al. 2013; Guerreiro et al. 2016)). Therefore, the *HoxD* TAD boundary is initially established in the absence of transcription within a ca. 50 kb window matching a large part of the *HoxD* cluster, likely in response to architectural proteins and/or other factors intrinsic to chromatin structure. Upon transcriptional activation, this border is refined and matches the transition between active and inactive *Hoxd* promoters. Consequently, the exact position of this boundary slightly varies along various cell types or tissues analyzed, in agreement with the proposal that insulation between TADs is favored by sharp transitions both in CTCF binding sites and in transcriptional activity (Zhan et al. 2017).

While the refinement of the boundary is associated with gene activity, the global positioning of the border at the *HoxD* cluster and its architecture may in turn cause a restriction in the subset of genes capable of responding to either TADs whenever they become activated. For instance, in both proximal limb bud cells and intestinal caecum where T-DOM is active, the border is established between *Hoxd11* (positive) and *Hoxd12* (negative) (Andrey et al. 2013; Delpretti et al. 2013). In the mammary gland however, this boundary seems to form between *Hoxd9* (positive) and *Hoxd10* (negative) (Schep et al. 2016). In contrast, when C-DOM is activated, either in distal limb cells or in the developing genitals, the boundary is found somewhere between *Hoxd10* (active) and *Hoxd9* (weakly active) (Lonfat et al. 2014). This partial overlap in the subsets of genes responding either to T-DOM or to C-DOM may reflect structural constraints and thus participate to the functional exclusivity observed at this locus thus far, for the two TADs are never activated concomitantly.

### Active *versus* inactive TADs and loop extrusion

Upon TAD functional activation, specific changes were observed in the interaction profiles, reflecting several states of configurations in chromatin architecture as reported earlier (Li et al. 2012; Sanyal et al. 2012; Berlivet et al. 2013; Rao et al. 2014; Dixon et al. 2015; Bonev et al. 2017). While some contacts were constitutive, others appeared only when the TAD enhancers were at work. For example, the 5’-located *Hoxd* genes are constitutively anchored to both island-1, which locates close to the next TAD border, as well as to island-2 and island-5. In distal cells however, where C-DOM shows high levels of H3K27ac, island-3 and other regions were also contacted and could thus be used as hallmarks of C-DOM transcriptional activity.

Our mapping of both CTCF sites and H3K27 acetylation suggest that once functionally active, enhancers within one TAD contact various subsets of target genes, depending on the cellular context. These distinct series of neighbor target genes are delimited by various combinations of bound CTCF sites, as if the presence of CTCF molecules would help define the different set of target genes responding in any given regulatory context. While the dynamic role of CTCF in marking chromatin domains has been documented (e.g. (Narendra et al. 2015)), we suggest here that series of bound CTCF sites in close proximity *in cis* may allow for tissue-specific interactions between long range enhancers and distinct contiguous groups of target *Hoxd* genes, perhaps through the selection of different CTCF sites in various contexts. However, the CTCF profiles analyzed in this study are invariable between distal and proximal limb cells and TAD borders tend to be co-occupied by CTCF and cohesin complexes (see (Ghirlando and Felsenfeld 2016), suggesting that other tissue-specific factors may be involved in the definition of sub-groups of target *Hoxd* genes, in combination with constitutive proteins. Our deletion analyses support this view, since the most notable effects on chromatin architecture were scored when the posterior part of the cluster was affected, i.e. the DNA interval where CTCF sites are concentrated. In such cases, deleting parts of the cluster would reconfigure the micro-architecture thus leading to another set of possible target genes.

Within the *HoxD* cluster, the CTCF sites located centromeric to *Hoxd11* are orientated towards C-DOM, whereas sites located telomeric point towards T-DOM (Fig. S7B). Also, the sites occupied by CTCF within either TADs and which correspond to the strongest interactions with *Hoxd* genes, including those at the two remote TAD boundaries, are mostly orientated towards the *HoxD* cluster. These observations support a loop extrusion model for the formation of these 3D chromatin domains (Rao et al. 2014; Fudenberg et al. 2016; Sanborn et al. 2015). In this view, the multiple copies of CTCF sites in *cis* around *Hoxd* genes may offer different possibilities for determining the extent of loop extrusion and thus lead to distinct positions of the boundary in various contexts, perhaps due to slightly different stabilization of loop-extruding factors (for instance cohesin) at neighboring but distinct sites.

However, while our mutant alleles can be generally reconciled with this interpretation, some alleles are more difficult to integrate into this model. The large majority of our deletion alleles indeed maintain at least one pair of CTCF sites with opposed orientations, which could thus account for the persistence of a *HoxD* boundary even with a much weaker insulation potential. For instance in the *HoxD^del(1-13)d9lac^* condition, while all native CTCF sites orientated towards T-DOM are deleted, two opposite sites are brought by the *Hoxd9* transgene, which may account for the weak boundary still observed. In contrast, the *HoxD^del(1-10)^* allele lost all CTCF sites orientated towards T-DOM and kept only those sites pointing towards C-DOM (Fig. S7B). Despite this imbalance in site orientation, the interactions observed with the T-DOM-specific bait CS38 revealed a strong insulation effect, virtually identical to that scored when the C-DOM-specific bait island-4 was used. This suggests that the series of CTCF sites orientated towards T-DOM at the position of the boundary are not pre-required to the formation of the telomeric TAD. In this case however, the centromeric TAD should not be affected (all appropriate CTCF sites remain) and this domain may prevent interactions with T-DOM region CS38 to occur.

The *HoxD^del(attP-Rel5)d9lac^* allele provided us with the minimal boundary elements potentially necessary for insulating the two TADs. A single *Hox* gene was left, with a transcriptionally active promoter fully capable to respond to both C-DOM and T-DOM enhancers. In addition, this transgene harbored two occupied CTCF sites with opposite orientations, each of them facing its neighboring regulatory landscape. However, no particular insulation effect was detected in this condition and the transgene responded rather correctly to all surrounding enhancers now belonging to a large and unified TAD (see below). This result suggests that at this particular locus, the required border between TADs is built through an additive effect of many elements, which altogether provide the tightness necessary to prevent illegitimate enhancer-promoter interactions. This may also explain why this boundary is still clearly detected in the almost complete absence of CTCF (data in (Nora et al. 2017))

### Attracting landscapes, tolerated interactions and border directionality

In several partial deletion alleles, ectopic interactions leaked over the border, leading to contacts between some *Hoxd* genes and the ‘wrong’ TAD. These leakages were not passive but instead often coincided with the activity of the TAD involved, as if an active TAD could attract ectopic contacts more efficiently than when inactive. Shared transcription factors and RNA polymerase II occupy both active enhancers and the set of target promoters, likely stabilizing the interaction (Kieffer-Kwon et al. 2013; Mousavi et al. 2013) and thus making trans-boundary contacts easier to detect by chromosome conformation capture. The role of cohesin and mediator, which also seem to be enriched according to the differentiation status (Phillips-Cremins et al. 2013) could also be investigated in this context.

Two different types of ectopic contacts were observed in our deletion alleles. The first one involved contacts between regulatory sequences belonging to both TADs such as for example increased interactions between the T-DOM CS38 sequence and regulatory islands located within C-DOM. While these contacts were scored, their deleterious effects are unlikely since they did not involve the mis-regulation of any important transcription unit. The second category of ectopic contacts involved the leakage of specific *Hoxd* target genes into another TAD, thus bringing them under the control of a distinct set of enhancers. For example some deletions allowed *Hoxd4* to contact C-DOM and thus be expressed in distal limb cells, whereas some others activated *Evx2* into proximal limb cells, due to its illegitimate interactions with T-DOM. In this case the mis-regulation of *Hoxd* genes could lead to potential alterations in morphological development. Accordingly, the tight and resilient boundary observed at the *HoxD* locus may have primarily evolved to prevent the precocious and ectopic expression of neighboring *Hoxd* genes during development, rather than to avoid inter-TADs contacts.

This possibility is supported by the apparent directionality in the leakage potential of flanking genes in control and deletion alleles. The analyses of several deletions indeed pointed to a general tendency for 3’-located genes (*Hoxd4; Hoxd8)* to respond to C-DOM enhancers more readily than 5’-located genes (*Evx2*; *Hoxd13*) would respond to T-DOM enhancers, as if the boundary effect was more efficient in blocking proximal than distal regulations to reach the opposite extremities of the gene cluster. This property could already be observed in control mice, with digit enhancers leaking up to *Hoxd9*, even though only *Hoxd13* showed an unambiguous function during digit development. In contrast, proximal enhancers are readily blocked at the *Hoxd11* locus, one of the key genes for zeugopod development (Davis et al. 1995) and no leakage in contact is observed onto *Hoxd13.*

### An adaptation to posterior prevalence

This directional property may be related to the rule of posterior prevalence, a functional property of posterior HOX proteins to often suppress the function of anterior ones when co-expressed, unlike in the opposite situation (Gonzalez-Reyes et al. 1990; Bachiller et al. 1994). As a consequence, the ectopic expression of group 13 *Hox* genes causes dramatic phenotypic alterations (Young et al. 2009)(see (Mallo et al. 2010)). Therefore, while contacts between digit enhancers and more 3’-located, anterior *Hoxd* genes may not have any functional consequences as long as *Hoxd13* is expressed there, the opposite situation where *Hoxd13* would respond to various T-DOM enhancers may readily elicit abnormal phenotypes. The sensitivity of this effect was previously observed when a subtle *Hoxd13* gain of function in proximal limb bud cells was enough to induce a light limb malformation (Tschopp and Duboule 2011). As a consequence, the *HoxD* TAD boundary must be very stringent in blocking proximal enhancers as a necessary adaptation to posterior prevalence, whereas digit enhancers may have interactions with various target *Hoxd* genes without any particular effect. While the mechanistic basis of this directionality is unclear, it may rely upon the complex distribution and various interaction strengths of architectural proteins at the boundary (see above), as reported in other cases (e.g. (Symmons et al. 2014; Tang et al. 2015; Ji et al. 2016).

### TADs ontology at the *HoxD* locus

*HoxD* lies between two TADs carrying distinct sets of regulatory sequences and operating one after the other, in an exclusive manner. While the necessity to functionally separating two sets of target genes is discussed above, the question remains as to whether groups of enhancer sequences with distinct specificities must segregate into different landscapes to properly work in space and time as previously suggested (Beccari et al. 2016). We show that in our largest deletion, a single TAD now forms containing at least five out of the seven digit regulatory elements as well as a strong proximal enhancer. Our targeted reporter transgene allowed us to conclude that most ‐if not all- enhancers could still exert their regulatory potential over this single promoter. This suggests that the two-TADs organization at the *HoxD* locus did not evolve to provide particular structural environments to series of holo-enhancers such as to optimize their regulatory inputs once they become functional. Instead, this partitioning in global regulations might be necessary to properly assign sub-sets of target *Hoxd* genes to their appropriate enhancers. This observation, added to a previous experiment showing that two enhancers located far from one another within the same TAD could work efficiently when associated into a unique small transgenic construct (Lonfat et al. 2014), supports a modular view of enhancer organization within TADs, whereby relative positions may not importantly impact upon their functionalities.

## MATERIALS AND METHODS

### Animal experimentation and mouse mutant lines

All experiments were performed in agreement with the Swiss law on animal protection under license number GE 81/14 (to D. D.). All tissues were obtained from E12.5 mouse embryos coming from the *HoxD^del(10-12)^*, *HoxD^del(9-12)^*, *HoxD^del(8-13)rXII^*, *HoxD^del(8-13)d11lac^*, *HoxD^del(1-10)^*, *HoxD^del(1-13)d9lac^*, *HoxD^del(1-13)d11lac^*, *HoxD^del(10-13)rXII^, HoxD^del(11-13)rXII^*, and *HoxD^Rel5d9lac^* mutant stocks, already reported by this laboratory. The *HoxD^del(attP-Rel5)d9lac^* was generated by TAMERE (Herault et al. 1998) between the *HoxD^attP^* (Andrey et al. 2013) and *HoxD^Rel5d9lac^* (Montavon et al. 2011) lines. To facilitate reading of the figures, the names of the alleles were reduced to the aforementioned superscripted annotations. All experiments were conducted using homozygous embryos derived from heterozygous crosses.

### Mutant genomes *in silico*

For the *HoxD^del(10-12)^*, *HoxD^del(9-12)^*, *HoxD^del(8-13)rXII^*, *HoxD^del(8-13)d11lac^*, *HoxD^del(1-10)^*, *HoxD^del(1-13)d9lac^*, *HoxD^del(1-13)d11lac^* and *HoxD^del(attP-Rel5)d9lac^* deletion lines, a corresponding mutant genome was built *in silico* to allow for a precise mapping of reads without apparent gaps. Chromosome 2 of these mutant genomes was built using mm10 as a backbone and applying the insertion/deletions using the package seqinr (Charif and Lobry 2007) in R software (Team 2008) (Supplemental Fig. S2).

### Hi-C

Distal and proximal forelimb and hindlimb bud tissue from either control, *HoxD^del(1-13)d9lac^* and *HoxD^del(attP-Rel5)d9lac^* were micro-dissected and collected individually. Cells were dissociated in 10% FBS/PBS with collagenase XI (C7657, SIGMA) to a final concentration of 0.4-0.6µg/µl, incubating samples for 60 minutes at 37°C in agitation (650 rpm). The cell suspension was strained and fixed for 10 minutes in formaldehyde (2% final concentration in 10% FBS/PBS). Cells were then centrifuged to discard the supernatant and frozen at −80°C until used subsequently, after genotyping. Hi-C libraries were generated using the HindIII enzyme as described in (Belton et al. 2012). Hi-C libraries were sequenced on an Illumina HiSeq4000 platform and 50 base paired end reads were obtained. Reads were mapped, filtered and bias corrected as described before (Lajoie et al. 2015; Giorgetti et al. 2016). The Hi-C datasets generated in this work as well as the mESC (Dixon et al. 2012) and CH12 (Rao et al. 2014) available datasets were processed identically. Briefly, read pairs were mapped independently, starting at 25bp and iterated every 5bp using bowtie2 (version 2.2.4 (Langmead et al. 2009)), as in (Imakaev et al. 2012) with parameter: ‐‐very-sensitive, either on mouse genome (mm10), or on the mutant genomes generated *in silico*. Each read was assigned to a fragment using the 5’ mapped-position shifted 3 bp toward the 3’ position to assign correctly the reads overlapping the cutting sites. The fragment assignment and mapping strand from R1 and R2 was combined and used to filter out the single-side mapped pairs, dangling-end pairs, error pairs and self-circle pairs. For each condition (same tissue and same genotype), two replicates were merged (fifteen replicates for CH12) and the interactions were filtered to discard duplicates. Each fragment was assigned to a bin (40 or 20kb) based on the position of the middle of the fragment and each valid interaction was assigned to a pair of bins (one bin for the R1 and one bin for the R2) and reported in the raw matrix. Prior to the ICE normalization (Imakaev et al. 2012) the rows and columns were masked if the sum of reads in this region was 10 fold less the expectation with uniform coverage, or if the number of fragments covered by at least two reads in this region was less than half the number of the fragments of this region. The ICEd matrixes were used for figures. In figures 1D and 6G, H, the difference between the two ICEd matrixes is plotted. All plots of matrixes were generated with R software (Team 2008). The insulation index in figure 6 was evaluated by TopDom (Shin et al. 2016) with a window size of 6 bins for the 40kb matrixes, corresponding to a −240kb, +240kb ‘diamond’. To call ‘consensus’ TADs out of which the TAD borders were called, TopDom algorithm was run with window sizes from 3 to 15 from the 40Kb binned matrices. Only the TADs present with the exact same coordinates in at least 40 % of the window sizes were considered as ‘consensus’. In supplemental Fig. S1, consensus TADs were called from 20Kb resolution for ESC and CH12 Hi-C data. To quantify the difference in contact intensities between the proximal and distal wild type datasets, a Wilcoxon rank sum test with continuity correction was performed on the ICEd values of every bin except the one on the diagonal in the C-DOM (chr2:73960000-74680000) and in the T-DOM (chr2:74720000-75600000), as called in the distal dataset. To be able to compare the contacts between the Hi-C data from *HoxD^del(attP-Rel5)d9lac^* and the Hi-C data from wild type, the mutant Hi-C data were mapped on the wild type mm10 genome. Before the ICEd normalization, the contacts involving bins representing deleted regions in the mutant genome (chr2:74400000-74760000) were removed from both wild type and mutant datasets. The computations were performed at the Vital-IT Center for high-performance computing of the Swiss Institute of Bioinformatics (http://www.vital-it.ch).

### 4C-sequencing

The distal and proximal parts of forelimb buds were dissected in cold PBS, placed in 250µl of PBS/10% FBS and digested in presence of collagenase XI (C7657, SIGMA) to a concentration of 0.4-0.6µg/µl. Samples were incubated at 37°C for 45 minutes in agitation. The cell suspension was strained through a mesh (352235, Falcon), fixed in 2% formaldehyde (in 10% FBS/PBS), lysed and centrifuged in order to obtain free nuclei precipitate, which were frozen at −80°C and stored. After genotyping, ten to fourteen pairs of each tissue were pooled in 500µl of 1.2x CutSmart Buffer (NEB, Ipswich, MA) and digested with *NlaIII* (NEB) as described in (Noordermeer et al. 2011). After the first 4h ligation, samples were digested using *DpnII* (NEB) in the corresponding buffer overnight and ligated again for 4h. Short fragments and nucleotides were discarded with the Nucleotide Removal Kit (QIAGEN, Germany) and libraries were prepared by means of 12 to 16 independent PCR reactions using 70 to 100ng of DNA on each (Supplemental Table S1). PCR products were pooled and purified using PCR purification kit (Qiagen). Up to twenty-two libraries were multiplexed either by combining different viewpoints or by means of 4bp barcodes added between the Illumina Solexa adapter sequences and the specific viewpoint inverse forward primer and sequenced using 100bp single reads on the Illumina HiSeq system. The obtained reads were de-multiplexed, mapped and analyzed using the pipeline present at BBCF HTSstation (http://htsstation.epfl.ch) (David et al. 2014) on the ENSEMBL mouse assembly GRCm38 (mm10). The profiles were smoothened using a window size of 11 fragments. The numbers of replicates obtained for each experiment are listed in Supplemental Table S2.

### 4C-seq normalization and quantifications

All the 4C-seq profiles were normalized to the distribution of reads along 5Mb upstream and downstream from each viewpoint region, except for LacZ viewpoint. The quantification of contact distribution along T-DOM (mm10, chr2: 74781516-75605516) and C-DOM (mm10, chr2: 73914154-74636454) was performed as read percentage of their reads sum, e.g. T-DOM/(T-DOM+C-DOM) x 100. The reads from the stated regions were obtained through the post-processing operations offered at HTSstation (http://htsstation.epfl.ch) (David et al. 2014). The quantification of contacts established at regulatory sequences was performed using the intersect BEDtools resource. The results show the distribution of 4C fragments at the given regions (mm10): island-1 (chr2:73970064-73983434), island-2 (chr2:74060473-74082287), island-3 (chr2:74177798-74223313), island-4 (chr2:74263814-74284643), island-5 (chr2:74289658-74313573), GCR (chr2:74445394-74498046), Prox (chr2:74604505-74639799), CS38-41 (chr2:75120051-75165771) and CS65 (chr2-75413472-75451553). Graphs and statistical analysis were performed with GraphPad Prism 7.

### 4C-seq relative cumulative frequency

For the relative cumulative frequency, the 4C-seq data were mapped to their respective newly generated genome and processed using the pipeline present at the BBCF HTSstation. The output used for the relative cumulative frequency was the segtofrag file. In the plot and for each dataset, the data was shifted in order to put the coordinates of the viewpoint at 0. Only the data between ‐1092537 and 2006380 for island-4 and between ‐1957157 and 1141528 for CS38 were used. These regions correspond to chr2: 73180041-76279897 in the wild type genome (mm10).

### DNA-FISH

3D DNA-FISH was performed as in (Morey et al. 2007; Fabre et al. 2015). Fosmids were used for both CS38-41 (WI1-2299-I7, mm10, chr2:75122702-75160145) and island-2 (WI1-109P4, mm10, chr2:74064904-74104783). Several BACs were used to cover T-DOM and C-DOM. T-DOM-1 (RPCI-23-190O13, mm10, chr2:74714710-74911321); T-DOM-2 (CH29-519G12, mm10, chr2:74893841-75119533); T-DOM-3 (CH29-617N10, mm10, chr2: 75131563-75340886), T-DOM-4 (CH29-6K11, mm10, chr2:75354051-75619849), C-DOM-1 (RP23-146O7, mm10, chr2:73821548-74029145); C-DOM-2 (RP23-427C9, mm10, chr2:74032726-74211877) and C-DOM-3 (RP24-222J8, mm10, chr2:74211948-74492098). For the *HoxD^del(8-13)rXII^* experiments, fosmids CS38-41 and island-2 were used, as well as CS38-41 and C-DOM-1, C-DOM-2 and C-DOM3. For *HoxD^del(attP-Rel5)d9lac^* (Supplemental Fig. S7), C-DOM-1, C-DOM-2, T-DOM-1, T-DOM-2, T-DOM-3, and T-DOM-4. Images were captured using an inverted Olympus IX81 microscope with a 60X Plan-Apo objective (numerical aperture of 1.42) and a B/W CCD ORCA ER B7W Hamamatsu camera. Stacks with a 200 nm step were saved as TIFF stacks, reconstructed and deconvoluted using FIJI (NIH, v1.47q) and Huygens Remote Manager (Scientific Volume Imaging). The distances between DNA-FISH signals were quantified using an automated spot and surface detection algorithm followed by visual verification and manual correction using IMARIS version 6.5, Bitplane AG and Matlab 7.5, MathWorks SA. Statistical significance analyses of distances were performed using the Kruskal-Wallis test followed by Dunn’s post test. The displayed representative images (Supplemental Figs. S6 and S7) were taken from distal forelimb samples.

### ChIP-seq

All H3K27ac, SMC1 and RAD21 experiments were processed as ChIP-seqs. All CTCF experiments were also processed as ChIP-seqs with the exception of *HoxD^del(8-3)rXII^*, where the ChIPmentation protocol was used (see below). Limb tissues were dissected and fixed in 1% formaldehyde/PBS for 10 minutes. Chromatin was sheared with either a tip-point sonicator (BioBlock Vibra-cell) or with a bath sonicator (Diagenode Bioruptor Pico) in order to obtain fragments ranging from 150 to 700bp. Chromatin was precipitated with anti-CTCF (61311, Active Motif), anti-RAD21 (ab992, Abcam), anti-SMC1 (A300-055A, Bethyl) or anti-H3K27ac (ab4729, Abcam) using agarose beads and following the Active Motif protocol. Libraries were done with at least 4ng of DNA following Illumina protocol and sequenced to 50bp single-end reads on Illumina HiSeq.

### ChIPmentation

Limb tissues were dissected, fixed and sonicated as for ChIP-seq experiments. CTCF ChIP for *HoxD^del(8-3)rXII^* was carried out using the ChIPmentation protocol of (Schmidl et al. 2015). Chromatin was incubated overnight with antibodies and magnetic beads were added for at least 3h afterwards. Washes were performed with TF-WBI, TF-WBIII and Tris-HCl 10mM pH8. Then, 1µl of transposase was added for 1 minute at 37°C and washes were repeated with TF-WBI and TET. qPCR was carried out to determine the amount of cycles to be applied during library amplification. Libraries were done using Nextera Custom adapter sequences and multiplexed for sequencing. All PCRs were done using KAPA PCR system (KM2605, KAPA Biosystems) after heating up the polymerase mix for 45 seconds. Library purification was performed with AMPureXP beads. A beads-to-sample ratio of 0.7:1 was applied to remove long fragments and the recovered supernatant was incubated in a beads-to-sample ratio of 2:1. Beads were then eluted using 25ul of Tris 10mM. Libraries were sequenced to 50bp-single read length on Illumina HiSeq.

### ChIP, ChIPmentation and RNA-seq analyses

The profiles of ChIP and ChIPmentation were obtained following this process: adapters and bad quality bases were removed with cutadapt (Martin 2011) version 1.8 options ‐m 15 ‐q 30 ‐a GATCGGAAGAGCACACGTCTGAACTCCAGTCAC for ChIP and ‐a CTGTCTCTTATACACATCTGACGCTGCCGACGA for ChIPmentation. Reads were further mapped using bowtie2 on the mm10 genome (Langmead and Salzberg 2012), version 2.2.4 default parameters. Bam files were merged for replicates. The coverage was obtained as the output of MACS2 (Zhang et al. 2008) version 2.1.1.20160309 command line: macs2 callpeak ‐t input.bam ‐‐call-summits ‐B. By default, MACS2 only kept one tag at the same location (the same coordinates and the same strand), which would remove all potential contaminants from 4C experiments. A summary of the ChIP/ChIPmentation analyses is given in supplemental Table S3. Motif orientation was assessed using the resources of the CTCFBSDB 2.0 database (http://insulatordb.uthsc.edu/) by focusing on the motifs identified as MIT_LM7 and their associated strands. For the CH12 lymphoblasts H3K27ac ChIP-seq, ENCODE files ENCFF001KBR and ENCFF001KBQ were analyzed the same way. The BAM files of CH12 lymphoblasts RNA-seq were downloaded from ENCODE ENCFF507RJZ and ENCFF469ZCH and merged. Only the uniquely mapped reads were kept for the coverage.

### RNA extraction and qPCR

Total RNA was extracted following QIAGEN’s RNEasy Minikit. RNA was retrotranscribed into cDNA using Promega GoScript Reverse Transcriptase. Custom SYBR probes were used for quantitative real time PCR (qPCR) in a Biorad CFX machine (96-well plates) or an ABIPrism machine (384-well plates). Fold inductions were assessed by the double delta-CT method being referred to *Tubb* expression levels. The primers used were those described in (Montavon et al. 2008) and (Delpretti et al. 2013). Graphs and statistical analysis were performed with GraphPad Prism 7.

### Beta-galactosidase staining and *in situ* hybridization

Embryos were fixed in 4% PFA/PBS for 30 minutes and washed for 10 minutes three times in PBS-T (0.1% Tween). Specimens were then stained at 37°C in a solution containing 5mM potassium hexacyanoferrate (III), 5mM potassium hexacyanoferrate (II) trihydrate, 2mM magnesium chloride, 0.01% sodium deoxycholate, 0.02% NP-40 and X-gal (1mg/ml) solution in PBS. After proper staining was achieved, the specimens were washed three times for 15 minutes in PBS-T, fixed again in 4% PFA/PBS (30 minutes) and washed again. Images were taken with a Leica MZFLIII microscope. Whole mount *in situ* hybridization were performed as described in (Woltering et al. 2009). Images were taken with a Leica MZFLIII microscope.

## ACKNOWLEDGEMENTS

We thank members of the Duboule laboratories for sharing material and discussions. We also thank Marion Leleu for help with the HTS station platform, Anamaria Necsulea for help with bioinformatics analyses, Christian Schmidl and Daan Noordermeer for help on ChIPmentation and 4C-seq experiments, respectively, Johan Gibcus and Jon-Matthew Belton for advices on Hi-C sampling and mapping as well as Bryan R. Lajoie for generating scripts for the analysis and Mylène Docquier and Brice Petit from the genomics platform in Geneva. This work was supported by funds from the Ecole Polytechnique Fédérale (Lausanne), the University of Geneva, the Swiss National Research Fund (No. 310030B_138662) and the European Research Council grants System*Hox* (No 232790) and Regul*Hox* (No 588029) (to D.D.), as well as the NIH grant HG003141 (to J.D.). J.D. is an investigator of the Howard Hughes Medical Institute.

## AUTHORS CONTRIBUTIONS

**ERC**: Planned the experiments, produced and analyzed the datasets and wrote the manuscript.

**LLD**: Bio-informatics analysis of the datasets, amended the manuscript.

**PJF**: Designed, produced and analyzed the DNA-FISH datasets, amended the manuscript.

**LB**: Produced some ChIP-seq datasets, amended the manuscript.

**IEI**: Produced some WISH datasets.

**THNH**: Produced mice and managed the mouse strains.

**YZ**: Performed Hi-C experiment.

**JD**: Planned the Hi-C analyses, co-wrote the corresponding section of the manuscript.

**DD**: Planned the experiments, analyzed datasets, wrote the manuscript.

## DATA REPOSITORY

All original and re-analyzed sequencing data have been deposited in the Gene Expression Omnibus (GEO). The study super-series can be found under the number GSE101717, which contains the subseries GSE101713 (4C-seq), GSE101714 (ChIP-seq) and GSE101715 (Hi-C).

